# Antigenic stimulation in conjunction with cytokine is required for mediating IL-17A production in human MAIT cells

**DOI:** 10.64898/2026.01.27.702041

**Authors:** Se-Jin Kim, Dylan Kain, Deborah A. Lewinsohn, Gwendolyn M. Swarbrick, Meghan E. Cansler, Benjamin N. Bimber, GW McElfresh, Emily B. Wong, Sharon Khuzwayo, Thomas Riffelmacher, David M. Lewinsohn

## Abstract

Mucosal-associated invariant T (MAIT) cells are donor unrestricted T cells capable of both antigen-specific adaptive responses and cytokine driven innate-like functions. Although human MAIT cells uniformly express *ROR*γ*t* and *IL23R*, they generally produce IFN-γ, and only a small fraction produces IL-17. Recent studies show that combined TCR and cytokine stimulation can elicit functional heterogeneity in blood-derived MAIT cells. Here, we investigate the role of IL-23/IL-23R signaling in mediating the function and transcriptional profiles of lung MAIT cell clones. We demonstrate that BAL-derived lung MAIT cell clones exhibit distinct cytokine profiles and variable *IL23R* expression. Short-term IL-23 stimulation triggers clone-specific transcriptional programs and *IL23R*-dependent upregulation of type 17-associated genes. Prolonged conditioning of lung MAIT cell clones with TCR (5-OP-RU) and cytokine (IL-23) stimulation induces stable IL-17A production along with unique transcriptional changes. TCR + IL-23 conditioning alone upregulates clone-specific and shared cytoskeletal/structural gene programs, whereas subsequent PMA/Ionomycin stimulation further induces IL-12 family signaling and metabolic genes. Together, these findings demonstrate that *IL23R* expression and TCR signaling are required for IL-17A production, highlighting that these conditions may be met in tissue environments where MR1-specific antigens and proinflammatory cytokines coexist.

## Introduction

Mucosal-associated invariant T (MAIT) cells are a family of T cells, highly prevalent in humans that can mediate antimicrobial defense. They are defined by their use of a semi-invariant T cell receptor (TCR) α-chain (*TRAV1-2-TRAJ33/TRAJ12/TRAJ20*), and by their dependence on histocompatibility complex (MHC) class I-related protein 1 (MR1), which allows MAIT cells to be categorized as donor unrestricted T cells^1^. MAIT cells can recognize small molecule metabolites derived from not only a broad array of microbial pathogens, but also from the host, drugs and drug-like molecules, as well as, synthetic and organic ligands^2–8^. MAIT cells represent 1-10% of circulating T cells in the peripheral blood, and are further enriched in airways and mucosal tissues such as the lung, liver, and gut^9–11^. As MAIT cells express a semi-invariant TCR α-chain, and rely on a nearly monomorphic presenting molecule, they were considered to be innate. However, through distinct TCR β-chain usage, MAIT cells can discriminate between microbial ligands and initiate pathogen-specific responses^12–14^. In addition to TCR-dependent activation, MAIT cells can also be directly activated by cytokines such as IL-12 and IL-18^15^. Both TCR-dependent and TCR-independent activation can lead to the production of proinflammatory cytokines such as TNF-α and IFN-γ, as well as cytotoxic molecules such as Granzyme B and Perforin to promote antimicrobial defense^16,17^. In certain mucosal tissues and during inflammatory disease states, MAIT cells preferentially produce IL-17A and IL-22 to maintain tissue homeostasis and regulate host-microbe interactions^18–20^. These combined features position MAIT cells as a unique T cell subset capable of both antigen-specific adaptive and cytokine driven innate-like effector functions.

Murine MAIT cells have distinct functional IFN-γ (MAIT1) and IL-17 (MAIT17) subsets, which can be distinguished by expression of T-bet and RORγt, respectively^21,22^. In humans, despite universal expression of RORγt, fewer than 5% of peripheral blood MAIT cells produce IL-17, while the majority produce IFN-γ^20,23–25^. This paradox is further compounded by the observation that tissue-resident MAIT cells are phenotypically distinct from blood MAIT cells and that MAIT cells can differentiate in response to inflammatory stimuli^17,21,25^. Garner *et al.* show that human MAIT cells in blood and liver are transcriptionally distinct, with liver MAIT cells enriched for activation and tissue-resident transcriptional profiles. Moreover, although human MAIT cells are transcriptionally homogeneous at rest, activation through TCR or cytokines (IL-12 and IL-18) induces stimulus-specific transcriptional and phenotypic profiles, such as expansion of IL-17-positive MAIT cells with elevated *IL17F* expression^17^. Recent studies further illustrate this functional plasticity in human blood MAIT cells exposed to distinct polarizing cues^26,27^. Wang *et al.* demonstrate that IL-17A-producing MAIT cell clones derived from peripheral blood mononuclear cells (PBMCs) gain IFN-γ-producing capacity upon stimulation with polarizing cytokines such as IL-12 or IL-23, whereas IFN-γ-producing clones maintain their original functional phenotype even with the same stimuli^26^. Together, these findings highlight the capacity of human MAIT cells to produce IL-17 through transcriptional and functional reprogramming and support the importance of tissue residency and inflammatory milieu in driving these changes.

While human lung MAIT cells express *RORC* and *IL23R* transcripts, IL-17 production has not been demonstrated^9,28^. For example, lung MAIT cells exhibit significantly higher expression of *IL23R, RORC,* and *IL17A* compared to non-MAIT CD8^+^ T cells^28^. However, fewer than 1% of human lung MAIT cells produce IL-17^9^. Building on recent studies demonstrating that dual TCR and cytokine stimulation drives functional heterogeneity in blood MAIT cells, we sought to determine whether IL-23/IL-23R signaling modulates the function and transcriptional profiles of lung MAIT cell clones. We used IL-23 as the polarizing cytokine since *IL23R* is highly expressed in both blood and lung MAIT cells. Here, we demonstrate that conditioning human PBMC-derived MAIT cells with TCR (5-OP-RU) and cytokine (IL-23) induces IL-17A production. In contrast, bronchoalveolar lavage (BAL)-derived lung MAIT cell clones exhibit distinct cytokine profiles and variable *IL23R* expression across lung MAIT cell clones. Short-term IL-23 stimulation of MAIT cell clones triggers *IL23R-*dependent upregulation of type 17-associated genes. Conditioning these clones with TCR + IL-23 results in unique transcriptional profiles and stable IL-17A production following PMA/Ionomycin stimulation. Together, these findings demonstrate that *IL23R* is required for lung MAIT cell clones to induce a specific network of signaling and metabolic genes that reinforce IL-17 production upon TCR + IL-23 conditioning and stimulation.

## Results

### Conditioning of Human MAIT cells with TCR and cytokine enables IL-17A production

To determine if conditioning with 5-OP-RU (TCR) and IL-23 can induce IL-17 production, we first established an *in vitro* conditioning model that was modified from a 5-OP-RU stimulation protocol previously published^29^. Specifically, we conditioned peripheral blood mononuclear cells (PBMCs) from a healthy donor with 5-OP-RU on Day 1 and 7 to induce TCR-dependent activation, with or without the addition of the polarizing cytokine IL-23 every other day (Figure 1A). To evaluate the functional capacity of these clones after conditioning, we stimulated them on Day 12 with PMA/Ionomycin and assessed cytokine production. MAIT cells, defined as CD8^+^TRAV1-2^+^5-OP-RU-tet^+^ cells, conditioned with 5-OP-RU in the absence of IL-23 produced only IFN-γ (Supp. Figure 1, Figure 1B, C). In contrast, conditioning with 5-OP-RU and IL-23 expanded both IFN-γ/IL-17A-producing and IL-17A only-producing MAIT cells. Notably, conditioning with TCR + IL-23 induced IL-17A production in a minor proportion of MAIT cells that were previously IFN-γ-only producers, suggesting that MAIT cells can acquire new effector functions in response to polarizing cytokines. These findings demonstrate that human blood MAIT cells can adopt an IL-17A-producing phenotype following IL-23 conditioning.

**Figure 1.**
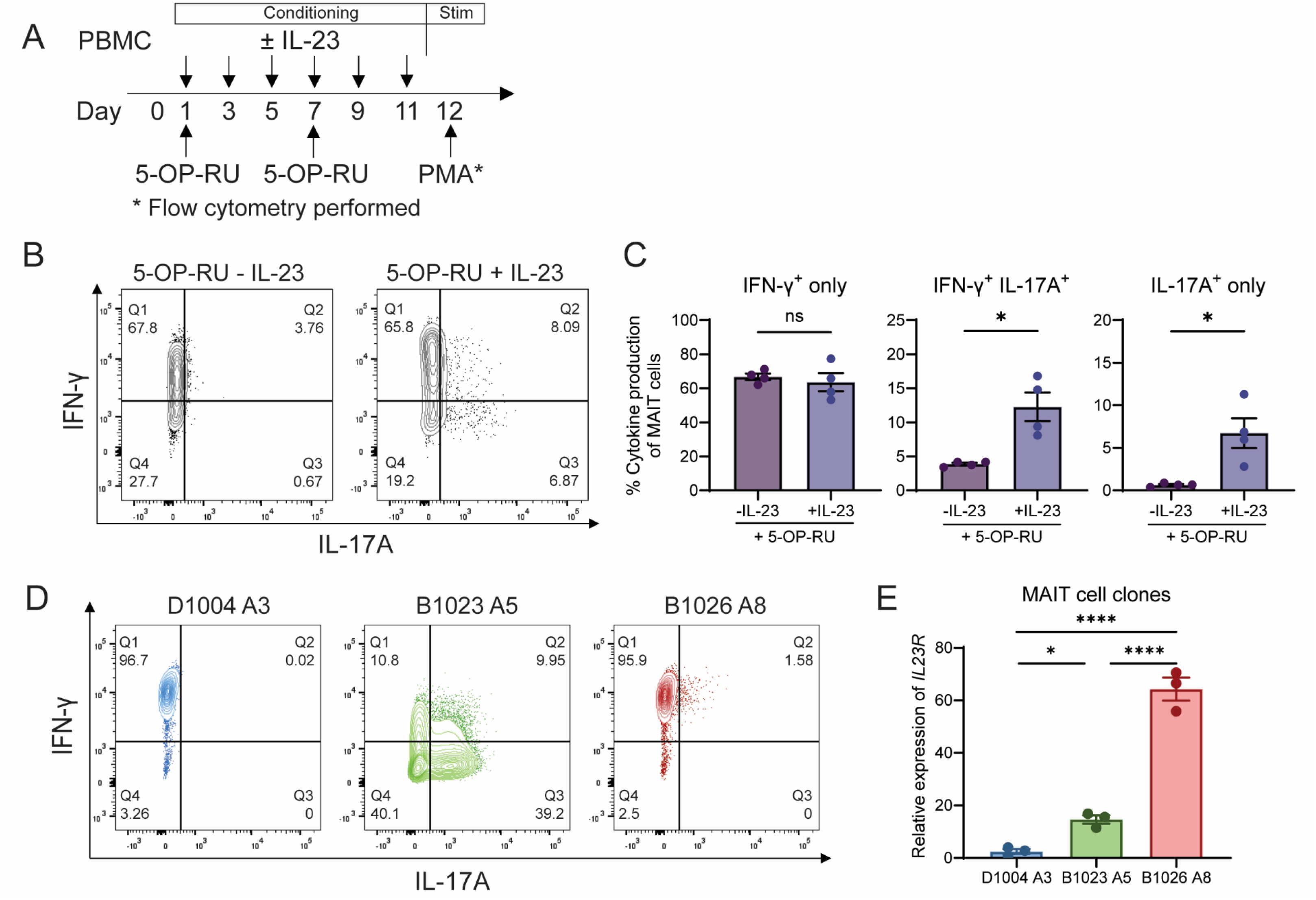
Human MAIT cells can produce IL-17A upon IL-23 stimulation and lung MAIT cell clones exhibit variable *IL23R* expression. (A) Experimental design: *in vitro* conditioning of PBMCs with 5-OP-RU and IL-23. Cells were incubated with PMA/Ionomycin for 3 hours on Day 12 for flow cytometry analysis. (B) Representative flow cytometry plot showing production of IFN-γ and IL-17A by MAIT cells conditioned with 5-OP-RU ± IL-23. MAIT cells are defined as live CD8^+^TRAV1-2^+^5-OP-RU-tet^+^ cells. (C) Percent cytokine production of IFN-γ^+^-only, IFN-γ^+^IL-17A^+^, and IL-17A^+^-only by MAIT cells. Data plotted as mean±SEM and pooled from four independent experiments. Two-tailed paired Student’s t-test. ns = not significant, * = p ≤ 0.05. (D) Representative flow cytometry plot showing production of IFN-γ and IL-17A by BAL-derived lung MAIT cell clones (D1004 A3, B1023 A5, B1026 A8) after incubation with PMA/Ionomycin for 3 hours. MAIT cells are defined as live CD8^+^TRAV1-2^+^5-OP-RU-tet^+^ cells. (E) Relative gene expression levels of *IL23R* compared to *GAPDH* in lung MAIT cell clones and normalized to lung non-MAIT CD8 T cell clone (D0033-JE1). Data plotted as mean±SEM and pooled from three independent experiments. Ordinary One-Way ANOVA Tukey’s multiple comparisons test. * = p ≤ 0.05, **** = p ≤ 0.0001.

### Human lung MAIT cell clones can produce IL-17 and exhibit variable IL23R expression

To further explore the functional consequences of IL-23 stimulation, we evaluated MAIT cell clones that were isolated from BAL samples collected from participants in South Africa^9^. As shown in our previous work, MAIT cell clones derived from lung express *IL23R*, but do not produce IL-17A^9,28^. Unexpectedly, we found discrete phenotypes with regards to IL-17A production. Clones D1004 A3 and B1026 A8 predominantly produced IFN-γ following PMA/Ionomycin stimulation. In contrast, the B1023 A5 clone produced both IFN-γ and IL-17A (Figure 1D). Given that we were now observing stable IL-17 production in a clone, it provided a unique opportunity to explore the mechanisms underlying IL-17A production. Since IL-23 promoted the functional conversion in blood MAIT cells, we examined *IL23R* expression in these lung MAIT cell clones. We found that clones D1004 A3 and B1026 A8 had low and high *IL23R* expression, respectively (Figure 1E). In contrast, B1023 A5, a clone with stable IL-17A production, expressed an intermediate level of *IL23R* (Figure 1E). The varying levels of *IL23R* expression and incomplete correlation between IL-23R levels and IL-17 production among these clones led us to investigate the mechanism of IL-23/IL-23R signaling and to identify additional gene programs that regulate IL-17A production.

### Increasing levels of IL-23R drives distinct expression profiles

To further assess the role of IL-23/IL-23R signaling, we conducted single-cell RNA-sequencing (scRNA-seq) on clones D1004 A3, B1023 A5, and B1026 A8, hereafter referred to as *IL23R*-low (A3), *IL23R*-med (A5), *and IL23R*-high (A8), that were unstimulated, stimulated with IL-23 overnight, or stimulated with PMA/Ionomycin for 3 hours. Stimulation with PMA/Ionomycin was chosen to ensure that each clone was viable and to confirm the IL-17 expression phenotype described above. The UMAP dimensionality reduction of each clone is shown in Figure 2A. Stimulation with PMA/Ionomycin for all three clones resulted in a separate cluster reflecting a distinct transcriptional profile as compared to unstimulated or IL-23 stimulated cells. In contrast, stimulation with IL-23 revealed three distinct patterns. *IL23R*-low (A3) clone did not result in a unique cluster following IL-23 stimulation. In contrast, both *IL23R*-med (A5) and *IL23R*-high (A8) clones responded to IL-23 stimulation, with more distinct clusters in A8 than A5. These results suggest that increasing levels of *IL23R* expression result in transcriptional profiles that were more distinct from unstimulated cells, which nonetheless did not correlate with the ability to produce IL-17 following PMA/Ionomycin stimulation.

**Figure 2.**
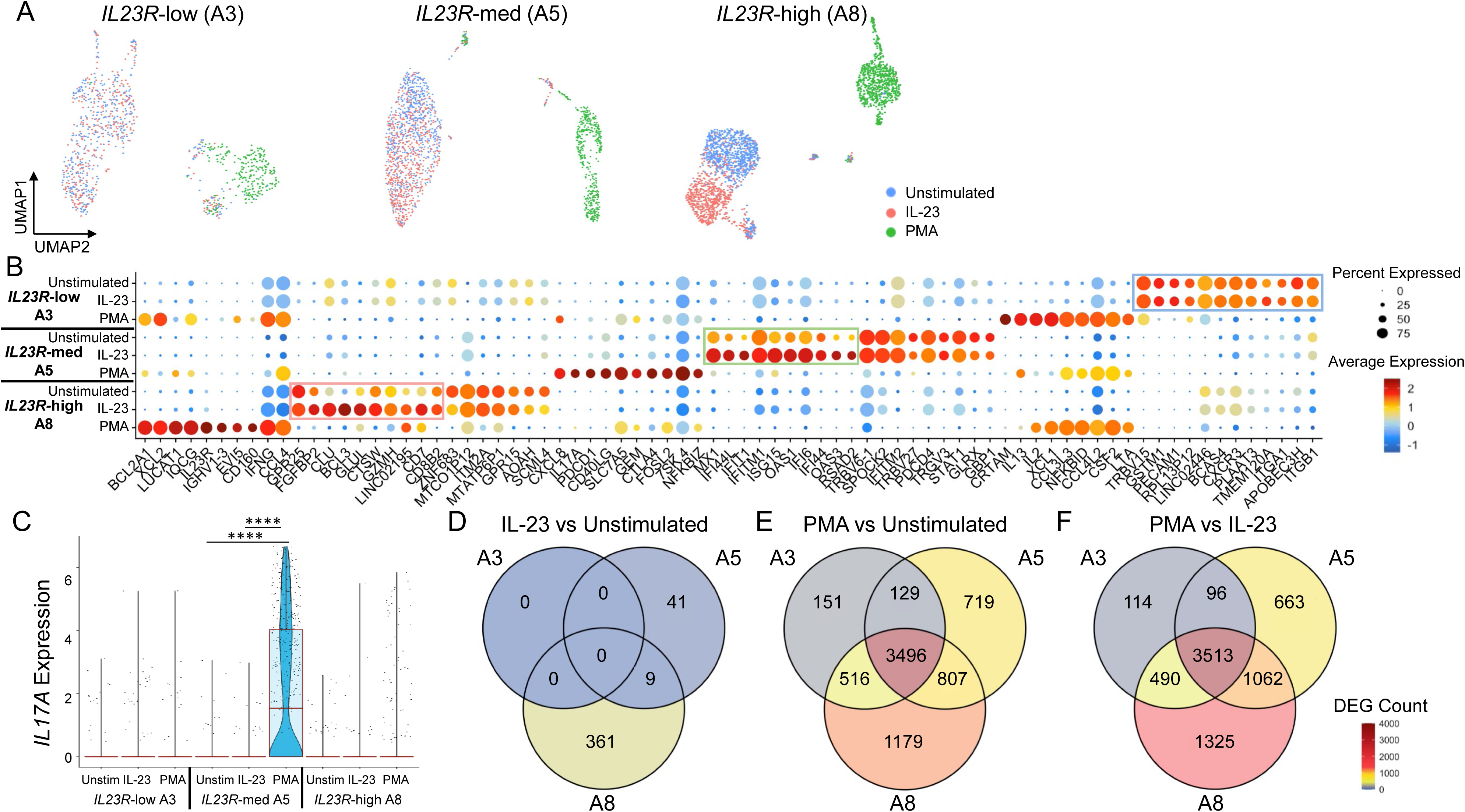
Increasing levels of *IL-23R* drives distinct expression profiles. (A) UMAP reductions of *IL23R*-low (A3), *IL23R*-med (A5), and *IL23R*-high (A8) for unstimulated (blue), IL-23 stimulation (red), or PMA/Ionomycin stimulation (green). (B) Dot plot of the of top differentially expressed genes defining each clone and stimulation state. Boxes indicate cluster of genes upregulated with short-term IL-23 stimulation for each clone. (C) *IL17A* transcript levels across clones with unstimulated, IL-23 stimulation, or PMA/Ionomycin stimulation. Kruskal–Wallis and post-hoc pairwise Wilcoxon with BH correction. **** = p ≤ 0.0001. (D–F) Venn diagrams showing numbers of differentially expressed genes (DEGs) for IL-23 versus Unstimulated (D), PMA/Ionomycin versus Unstimulated (E), and PMA/Ionomycin versus IL-23 (F).

To further explore the transcriptional profile induced by short-term IL-23 stimulation, we explored the top 10 genes differentially expressed by each clone following stimulation with IL-23 or PMA/Ionomycin (Figure 2B). In all three clones, stimulation with PMA/Ionomycin induced broad and robust transcriptional activation, marked by high expression of effector genes (*IL2*, *CSF2*, and *IFNG*), chemokines (*XCL1*, *XCL2*, *CCL3L3*, *CCL4L2*, and *CCL4*), and activation-associated genes (*CRTAM*, *PDCD1*, and *CTLA4*). Notably, PMA/Ionomycin stimulation induced *IL17A* expression in *IL23R*-med (A5) clone, corresponding to its ability to produce IL-17A upon activation, but there was no significant *IL17A* expression in the other clones (Figure 2C). Stimulation with IL-23 elicited no differentially expressed genes in the *IL23R*-low (A3) clone. In contrast, IL-23 triggered a selective, clone-specific transcriptional programs in the *IL23R*-med (A5) and *IL23R*-high (A8) clones. Specifically, the *IL23R*-med clone upregulated interferon-stimulated genes (*MX1*, *IFI44L*, *IFIT1*, *IFITM1*, *ISG15*, *OAS1*, *IFI6, IFI44*, *OAS3*, and *RSAD2*), whereas the *IL23R*-high clone upregulated genes such as *GPR25*, *FGFBP2*, *CLU*, *BCL3*, *GLUL*, *CTSW*, *GZMH*, *LINC02195*, and *CD8B2*, with *BCL3* and *CD7* representing type 17-associated genes^17,30^. These data elucidate the distinct patterns seen in the UMAP analysis.

To determine both shared and unique differential gene expressions (DEGs), we generated Venn-diagrams to compare clones in response to different stimulations (Figure 2D-F, Supp. Table 1 and 2). IL-23 stimulation induced transcriptional changes only in the *IL23R*-med (A5) and *IL23R*-high (A8) clones. The *IL23R*-high clone displayed the largest response (361 unique DEGs) and the *IL23R*-med clone showed a modest response (41 DEGs), while the *IL23R*-low (A3) clone exhibited no IL-23–responsive genes (Figure 2D, Supp. Table 1 and 2). In contrast, PMA/Ionomycin stimulation elicited broad transcriptional activation across all clones, with thousands of DEGs and a substantial shared core of 3,496 genes, alongside clone-specific signatures ranging from 151 DEGs unique to A3 to 1,179 DEGs unique to A8 (Figure 2E, Supp. Table 1 and 2). When directly comparing PMA/Ionomycin to IL-23 stimulation, the three clones again shared a large common set of 3,513 DEGs, with additional clone-restricted gene programs, most pronounced in A8 (1,325 unique DEGs) (Figure 2F, Supp. Table 1 and 2). These findings highlight the strong, largely overlapping transcriptional program induced by PMA/Ionomycin across MAIT cell clones, contrasted with the more selective and IL-23R-dependent response to IL-23.

Given the observed changes in type 17–associated genes, we next applied MAIT1 and MAIT17 transcriptional scores from Garner *et al.* to assess whether IL-23 or PMA/Ionomycin stimulation shifted the MAIT cell clones toward a MAIT1 or MAIT17 phenotype^17^. Across all three clones, PMA/Ionomycin stimulation exhibited a pronounced increase in MAIT1 scores, whereas short-term IL-23 stimulation had little effect on the MAIT1 score (Supp. Figure 2A). PMA/Ionomycin stimulation consistently reduced MAIT17 scores in all clones, regardless of *IL23R* expression (Supp. Figure 2B). IL-23 stimulation did not alter MAIT17 scores in the *IL23R*-low (A3) or in *IL23R-*med (A5) clones. However, in *IL23R*-high (A8) clone, IL-23 stimulation increased the MAIT17 score. Together, these findings indicate that PMA/Ionomycin strongly skews MAIT cells toward a MAIT1-like transcriptional profile while suppressing MAIT17 features, whereas IL-23 selectively enhances MAIT17 polarization in the *IL23R*-high clone.

### Conditioning of MAIT cell clones with 5-OP-RU in the presence of IL-23 results in the stable IL-17A production

As shown in Figure 2C, short-term exposure to IL-23 did not result in the upregulation of *IL17A* expression. Previous studies and Figure 1A-C show that prolonged TCR activation in combination with cytokine stimulation is required to elicit upregulation of IL17-associated genes. For example, Garner *et al*. demonstrated that combined TCR and cytokine stimulation for at least 68 hours strongly induces distinct *IL17F* upregulation, while Cole *et al.* showed that 3 days of TCR activation followed by PMA/Ionomycin stimulation enhances both IL-17A and IL-17F production^17,25^. Therefore, we modified our rapid expansion protocol to an experimental design using 5-OP-RU (TCR) and IL-23 (Cytokine) co-stimulation to more effectively assess the functional capacity and role of IL-23 on lung MAIT cell clones. Specifically, our conditioning protocol entailed co-culturing individual MAIT cell clones with irradiated LCLs and PBMCs, stimulating them with 5-OP-RU on Day 0 and Day 6, and adding IL-23 every other day where indicated (Figure 3A). To determine the functional profile of conditioned MAIT cells (defined as CD8^+^TRAV1-2^+^5-OP-RU-tet^+^ cells), cultured in the presence or absence of IL-23, we stimulated with PMA/Ionomycin and measured IL-17 and IFN-γ production (Figure 3A, Supp. Figure 3). As expected, IL-23 did not alter the phenotype of the *IL23R*-low (A3) clone (Figure 3B). In contrast, the *IL23R*-med (A5) clone showed a significant increase in IL-17A production when conditioned with IL-23 (Figure 3B, C). The *IL23R*-high (A8) clone acquired the capacity to co-produce IL-17A in conjunction with IFN-γ following conditioning with IL-23 (Figure 3B, C). These findings closely parallel the functional conversion we observed in human blood MAIT cells and underscore the functional adaptability of lung MAIT cell clones in response to polarizing cytokines.

**Figure 3.**
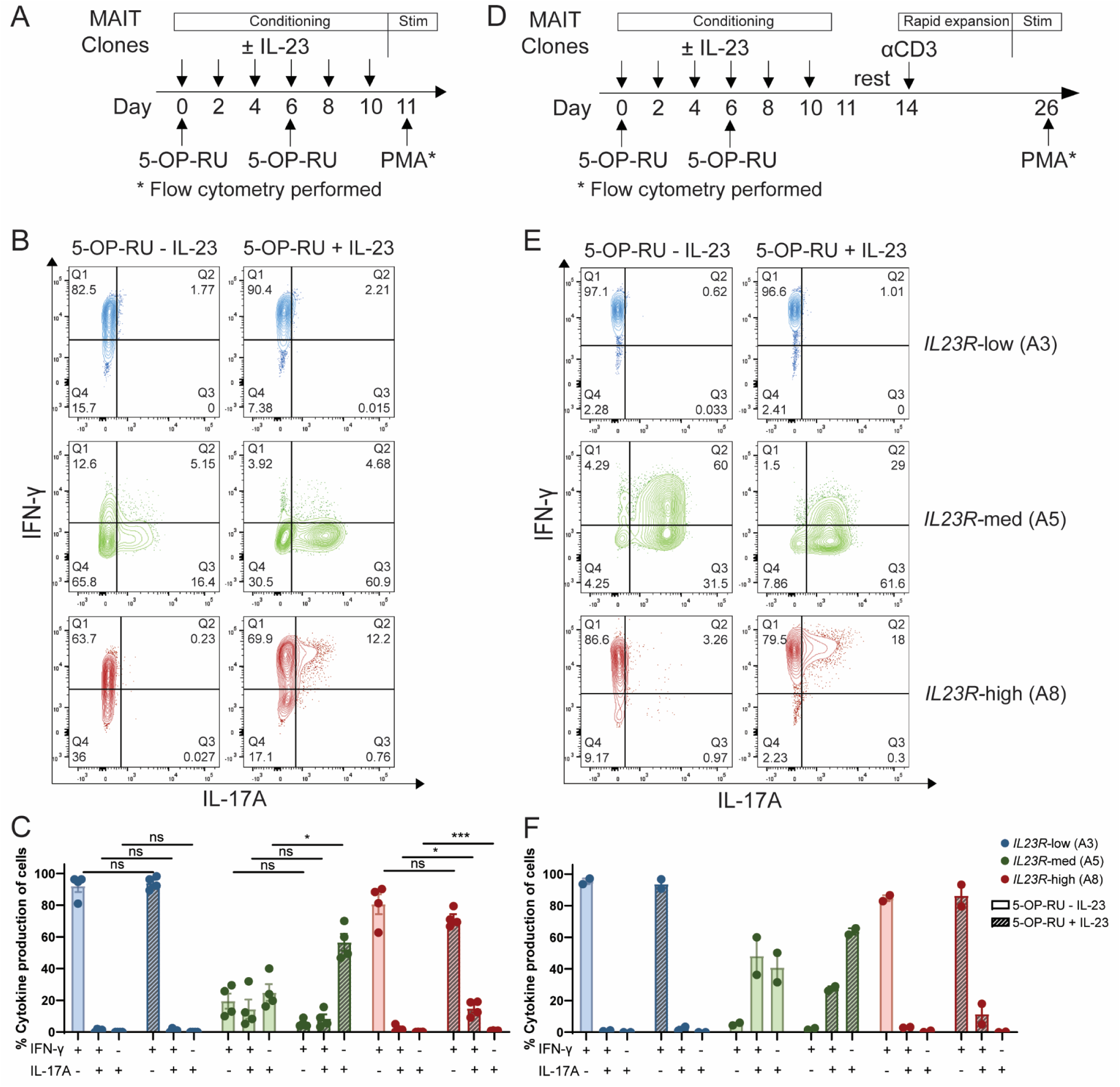
Conditioning of MAIT cell clones with 5-OP-RU in the presence of IL-23 results in the stable IL-17A production. (A) Experimental design: *in vitro* conditioning of lung MAIT cell clones with 5-OP-RU and IL-23. Cells were stimulated with PMA/Ionomycin for 3 hours on Day 11 for flow cytometry analysis. (B) Representative flow cytometry plot showing production of IFN-γ and IL-17A by lung MAIT cell clones conditioned with 5-OP-RU ± IL-23. MAIT cells are defined as live CD8^+^TRAV1-2^+^5-OP-RU-tet^+^ cells. (C) Percent cytokine production of IFN-γ^+^-only, IFN-γ^+^IL-17A^+^, and IL-17A^+^-only by MAIT cell clones. Data plotted as mean±SEM and pooled from four independent experiments. Two-tailed paired Student’s t-test comparing 5-OP-RU-IL-23 versus 5-OP-RU+IL-23 for each clone with cytokine produced. ns = not significant, * = p ≤ 0.05, *** = p ≤ 0.001. (D) Experimental design: Rapid T cell expansion after *in vitro* conditioning of lung MAIT cell clones with 5-OP-RU and IL-23. On Day 11, cells were rested for 3 days in low-dose IL-2 before the rapid T cell expansion for another 12 days. On Day 26, cells were stimulated with PMA/Ionomycin for 3 hours for flow cytometry analysis. (E) Representative flow cytometry plot showing production of IFN-γ and IL-17A by lung MAIT cell clones that were previously conditioned with or without IL-23 and then rapidly expanded. MAIT cells are defined as live CD8^+^TRAV1-2^+^5-OP-RU-tet^+^ cells. (F) Percent cytokine production of IFN-γ^+^-only, IFN-γ^+^IL-17A^+^, and IL-17A^+^-only by MAIT cell clones that were previously conditioned with or without IL-23 and then rapidly expanded. Data plotted as mean±SEM and pooled from two independent experiments.

To evaluate whether this phenotype is stable over time, we expanded these MAIT cell clones again in the absence of the polarizing cytokine IL-23. Specifically, after resting MAIT cell clones for three days in low-dose IL-2, we expanded the clones with TCR stimulation (αCD3) and co-stimulation (LCLs and PBMCs)^31^. At the end of this expansion, we measured cytokine production of lung MAIT cell clones following PMA/Ionomycin stimulation (Figure 3D). The resulting cytokine profiles remained consistent with those observed immediately following the initial 5-OP-RU and IL-23 conditioning phase (Figure 3B-C, E-F). The *IL23R*-low (A3) clone continued to produce only IFN-γ, regardless of prior exposure to IL-23. The *IL23R*-med (A5) clone, previously conditioned with IL-23, continued to show a trend toward increased IL-17A-only producing cells. Furthermore, the *IL23R*-high (A8) clone appeared to maintain its capacity to co-produce IFN-γ and IL-17A even in the absence of continued IL-23 exposure. Together, these results demonstrate that human lung MAIT cell clones undergo stable functional reprogramming in response to prolonged TCR and IL-23 co-stimulation.

### Single-cell profiling of human lung MAIT cell clones conditioned with 5-OP-RU and IL-23

To explore the mechanisms by which IL-23 conditioning led to the stable production of IL-17A, we performed scRNA-seq on the MAIT cell clones that were conditioned with 5-OP-RU ± IL-23 and then stimulated or not with PMA/Ionomycin (Figure 4A). As shown in Figure 4B, IFN-γ and IL-17A production were consistent with our earlier observations. Both the *IL23R*-med (A5) and *IL23R*-high (A8) clones increased IL-17A secretion without changes in IFN-γ when stimulated with PMA/Ionomycin after IL-23 conditioning (Figure 4B). Single-cell profiling of these clones also confirmed the protein expression as demonstrated by *IL17A* mRNA expression (Figure 4C). Both the *IL23R*-med (A5) and *IL23R*-high (A8) clones showed modest upregulation of *IL17A* expression with IL-23 conditioning, which was enhanced by PMA/Ionomycin stimulation.

**Figure 4.**
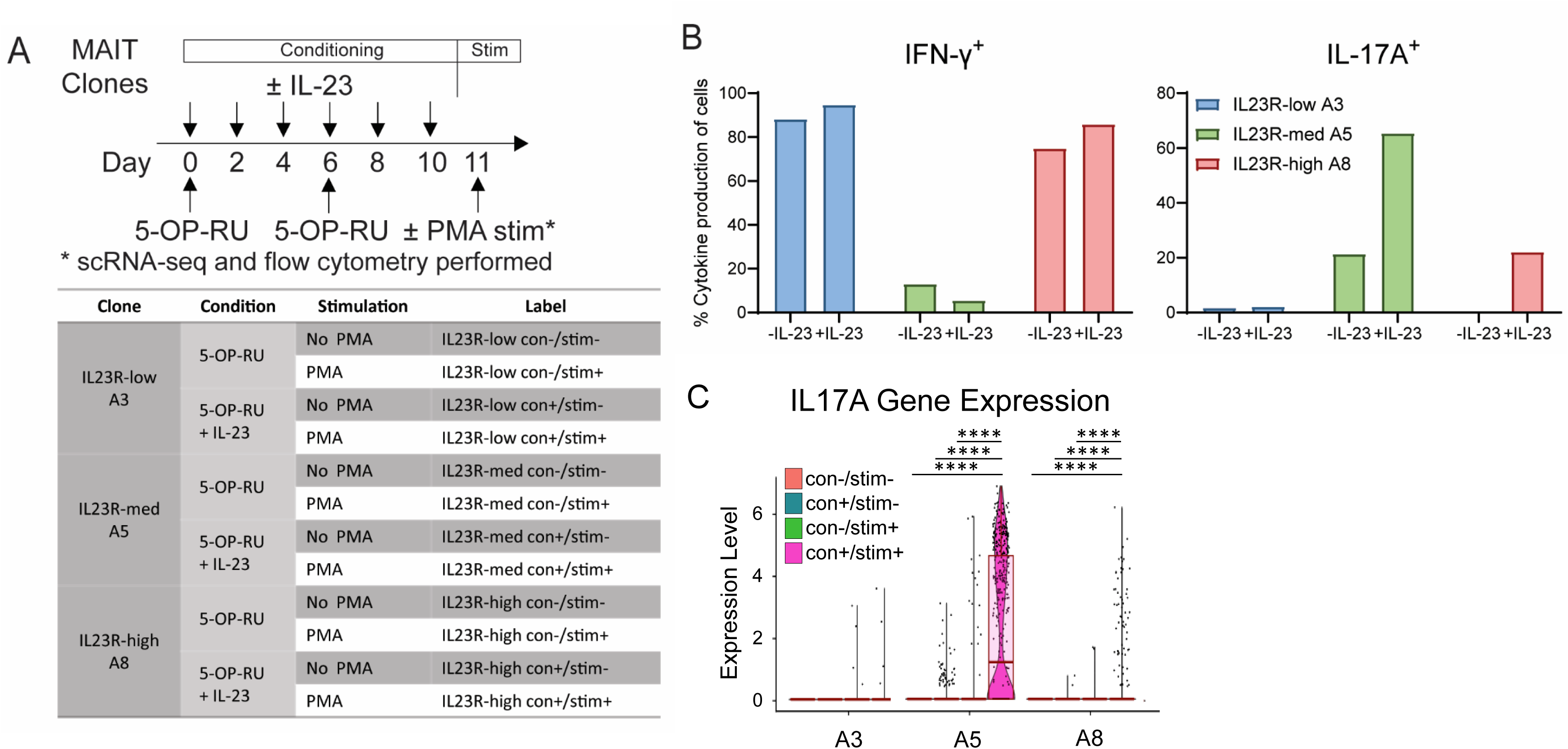
Functional phenotypes of human lung MAIT cell clones with IL-23 conditioning and PMA/Ionomycin stimulation. (A) Experimental design: *in vitro* conditioning of lung MAIT cell clones with 5-OP-RU ± IL-23. Cells were either stimulated or not with PMA/Ionomycin for 3 hours on Day 11 for flow cytometry analysis and single-cell RNA-sequencing (scRNA-seq). Labeling of each clone depending on conditioning and stimulation. (B) Validation of cytokine production after conditioning and stimulation following the experimental design in Figure 4A. Percent cytokine production of IFN-γ^+^-only and IL-17A^+^-only by MAIT cell clones after incubation with PMA/Ionomycin for 3 hours for flow cytometry analysis. (C) *IL17A* transcript levels across clones conditioned with 5-OP-RU ± IL-23 and then stimulated or not with PMA/Ionomycin. Kruskal–Wallis and post-hoc pairwise Wilcoxon with BH correction. **** = p ≤ 0.0001.

We next examined the dimensionality reduction for each conditioning regimen with subsequent stimulation (Figure 5A). The combination of TCR + IL-23 conditioning in both stimulated or not with PMA/Ionomycin showed no distinct clustering in the *IL23R*-low (A3) clone, whereas the *IL23R*-med (A5) and *IL23R*-high (A8) clones displayed partial and discrete clustering, respectively (Figure 5A). These data, therefore, demonstrate that TCR + IL-23 conditioning can alter the gene expression profile in the IL-23R expressing clones and that the degree of unique transcriptional phenotypes correlated with *IL23R* expression level. To further assess the genes differentially expressed after TCR and IL-23 conditioning from these UMAP clusters, we generated volcano plots and heatmaps comparing con-/stim- versus con+/stim- for each clone. In the *IL23R*-low (A3) clone, reactome pathway enrichment analysis of genes significantly downregulated following IL-23 conditioning revealed suppression of pathways associated with “interferon stimulated genes” and “adhesion and trafficking” (Supp. Figure 4A, D). In contrast, IL-23 conditioning significantly upregulated genes in both the *IL23R*-med (A5) and *IL23R*-high (A8) clones (Supp. Figure 4B, C). Although these two clones shared a subset of genes associated with “Cytoskeletal/Structure,” most upregulated genes were clone-specific, such as genes categorized as “Adhesion and Migration” for the *IL23R*-med (A5) clone and “Metabolic Activation” and “Lipid/Cholesterol Handling” for the *IL23R*-high (A8) clone (Figure 5B, C).

**Figure 5.**
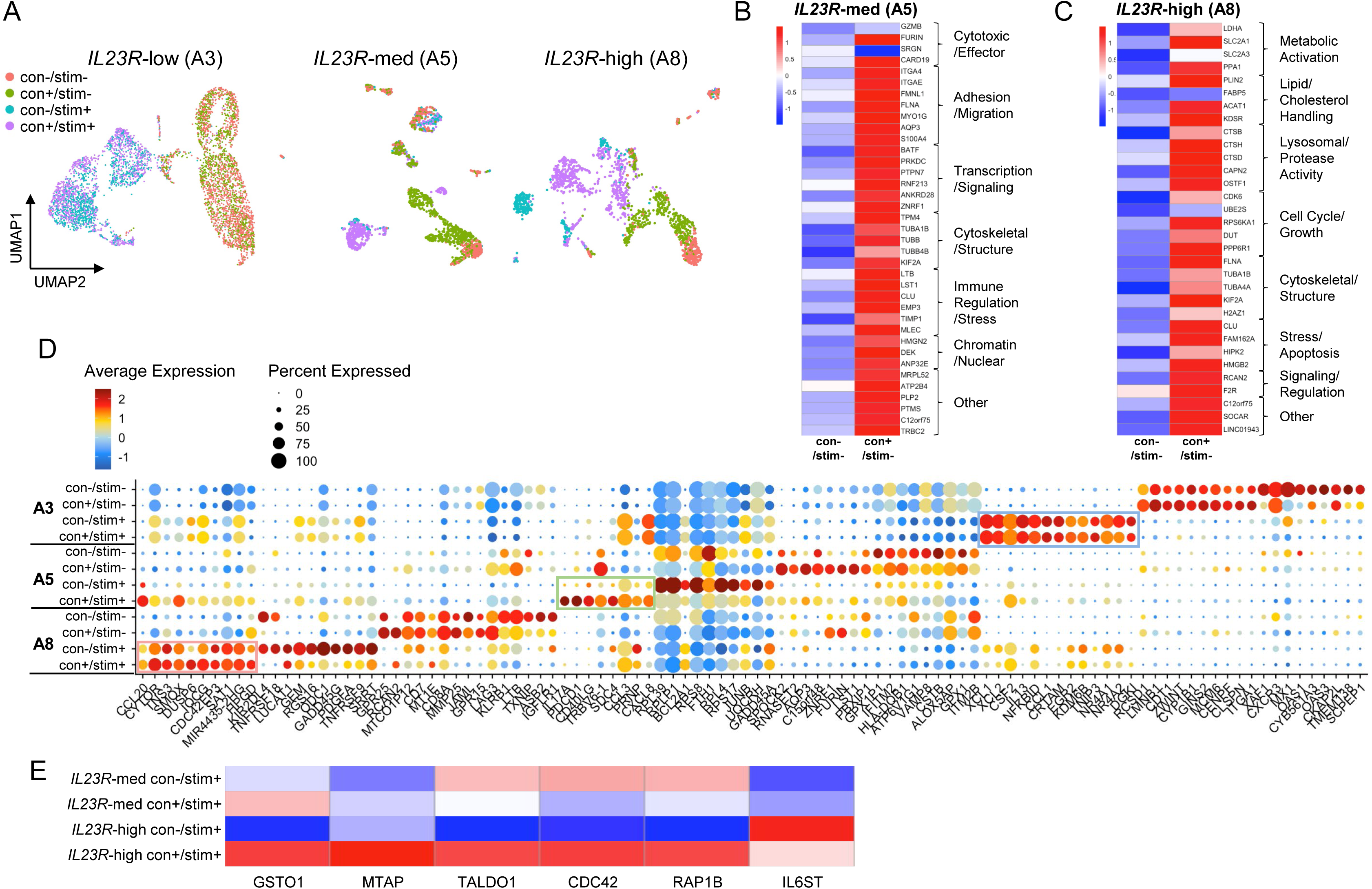
Single-cell transcriptional profiling of human lung MAIT cell clones conditioned with 5-OP-RU and IL-23. (A) UMAP projections of *IL23R*-low (A3), *IL23R*-med (A5), and *IL23R*-high (A8) clones under con-/stim-, con+/stim-, con-/stim+, and con+/stim+. (B) Heatmap of genes significantly upregulated in *IL23R*-med (A5) with IL-23 conditioning. Genes are grouped into shared categories and described. (C) Heatmap of genes significantly upregulated in *IL23R*-high (A8) with IL-23 conditioning. Genes are grouped into shared categories and described. (D) Dot plot of top differentially expressed genes per each clone with IL-23 conditioning and PMA/Ionomycin stimulation. Boxes indicate cluster of genes upregulated with IL-23 conditioning with PMA/Ionomycin stimulation for each clone. (E) Heatmap of specific genes significantly upregulated in *IL23R*-high (A8) with IL-23 conditioning and PMA/Ionomycin stimulation as part of IL-12 family signaling pathway enrichment analysis.

To further examine the changes in transcriptional profile with TCR + IL-23 conditioning and subsequent PMA/Ionomycin stimulation, we examined the top differentially expressed genes for each conditioning regimen (Figure 5D). We observed minimal transcriptional changes in the *IL23R*-low D1004 A3 clone, whereas both *IL23R*-med (A5) and *IL23R*-high (A8) clones upregulated unique gene sets in response to IL-23 conditioning as well as subsequent PMA/Ionomycin stimulation. Across all three clones, PMA/Ionomycin stimulation broadly activated effector molecules such as *CSF2* and *CD40LG*, as well as transcription factors including *NFKBID*, *EGR2*, *NR4A1* and *NR4A2*. In both *IL23R*-med and *IL23R*-high clones conditioned with TCR + IL-23, PMA/Ionomycin stimulation suppressed MAIT1-associated genes such as *CXCR3*, *XCL1*, and *XCL2*, as opposed to the upregulation of MAIT17-associated genes, including *IL17A* and *CCL20*^17^.

Given these transcriptional changes shown in Figure 5D, we next performed pathway enrichment analysis to understand the distinct biological processes associated with TCR + IL-23 conditioning and subsequent PMA/Ionomycin stimulation (Supp. Figure 5A-D). The *IL23R*-low (A3) clone showed no differentially expressed genes with conditioning and stimulation. In contrast, IL-23 conditioning alone for both the *IL23R*-med (A5) and *IL23R*-high (A8) clones was associated with cell cycle–related pathways, including “M phase,” “Mitotic anaphase,” and “RHO GTPase effectors” (Supp. Figure 5A, B), suggesting that the TCR + IL-23 conditioning regimen promotes cell proliferation and activation-related pathways. We next evaluated the resulting phenotype following PMA/Ionomycin stimulation. Following TCR + IL-23 conditioning and subsequent PMA/Ionomycin stimulation, the *IL23R*-med (A5) and *IL23R*-high (A8) clones exhibited different pathway enrichments. The *IL23R*-med clone was enriched for interleukin signaling, TCR signaling, and NF-κB–related pathways (Supp. Figure 5C), whereas the *IL23R*-high clone uniquely upregulated IL-12 family signaling, nucleotide metabolism-related genes, and reactive oxygen species detoxification (Supp. Figure 5D).

To further identify specific genes that drives IL-23 responsiveness, we generated a heat map of “IL-12 family signaling” from the pathway enrichment analysis and compared *IL23R*-med (A5) and *IL23R*-high (A8) clones (Supp. Figure 6). We focused on “IL-12 family signaling” pathway as IL-23 belongs to this cytokine family and shares components such as p40 subunit, *IL12RB1* receptor, and downstream JAK-STAT activation. In both clones, IL-23 induced genes implicated in IL-23 responsiveness and IL-17 secretion such as *IL12RB1* and *MIF*^32,33^. Interestingly, the *IL23R*-high (A8) clone uniquely expressed *IL6ST* and upregulated *RAP1B* and *CDC42* upon IL-23 conditioning, which are genes associated with STAT3 signaling and Th17 differentiation (Figure 5E)^34–36^. Moreover, the *IL23R*-high (A8) clone further upregulated metabolic/redox regulators such as *TALDO1*, *MTAP*, and *GSTO1* compared to the *IL23R*-med (A5) clone. Together, these findings reveal a network of signaling and metabolic genes that allow the *IL23R*-high (A8) clone to co-produce IFN-γ and IL-17A. Elevated *IL23R* expression appears to amplify STAT3-driven Th17 pathways and metabolic and redox programs, sustaining IL-17 production in response to IL-23 conditioning.

## Discussion

In humans, MAIT cells are largely characterized by their production of IFN-γ. Paradoxically, MAIT cells in both blood and lung express RORγt and IL-23R, raising the question of why IL-17A-producing MAIT cells have been so difficult to identify. Recently, two studies address this paradox by demonstrating that dual TCR and cytokine stimulation can skew the transcriptional and cytokine profiles toward IL-17A production^17,26^. In this study, we extend these observations by developing a conditioning protocol that combines TCR (5-OP-RU) with cytokine (IL-23) stimulation. We focused on IL-23 based on its receptor expression on both blood- and lung-derived MAIT cells. When we applied the conditioning regimen to blood-resident MAIT cells, this resulted in the production of IL-17A. Moreover, we identified lung-derived MAIT cell clones that showed heterogeneity in *IL23R* expression and IL-17A production. Following overnight IL-23 stimulation, we found that IL-23 responsiveness in lung MAIT cell clones was associated with the level of *IL23R* expression. We then applied the conditioning regimen, together with subsequent PMA/Ionomycin stimulation, and found stable IL-17A production and unique transcriptional changes in *IL23R*-expressing MAIT cell clones. These findings highlight that the environment can actively shape MAIT cell function and, to some extent, their functional plasticity.

To understand the impact of conditioning regimen on lung MAIT cell clones with variable *IL23R* expression, scRNA-seq revealed key transcriptional changes. For both *IL23R*-med A5 and *IL23R*-high A8 clones, IL-23 conditioning upregulated clone-specific genes, while also inducing a shared subset of genes associated with “Cytoskeletal/Structure.” This potentially suggests that IL-23 conditioning primes MAIT cells for activation and motility to facilitate their effector functions. Notably, the *IL23R*-med A5 – a clone with stable IL-17A production – upregulated “Adhesion/Migration” genes, such as *AQP3* and *S100A4*, key components of the MAIT17 gene signatures^17^. In contrast, the *IL23R*-high A8 clone induced “Metabolic activation” genes, such as *LDHA*, a glycolytic enzyme associated with RORγt expression in MAIT17 cells^37^. These findings support the idea that TCR + IL-23 conditioning biases human lung MAIT cells toward a MAIT17 transcriptional program and IL-17A effector function.

Pathway enrichment analysis comparing *IL23R*-med A5 and *IL23R*-high A8 clones elucidated transcriptional changes that enabled the *IL23R*-high A8 clone to acquire the capacity to co-produce IL-17A and IFN-γ following conditioning and stimulation. IL-23 conditioning selectively upregulated IL-12 family signaling in *IL23R*-high clone – a pathway that shares JAK/STAT activation via shared IL12RB1 receptor – and induced genes linked to STAT3 signaling and Th17 differentiation (*IL6ST*, *RAP1B*, *CDC42*), as well as metabolic/redox regulators (*TALDO1*, *MTAP*, *GSTO1*)^34–36^. Together, these findings provide a potential explanation for the observed functional plasticity: high *IL23R* expression potentiates IL-23/IL-23R signaling, engages STAT3-dependent and metabolic programs, and thereby promotes IL-17A production. These data would also argue that the presence of IL-23 in the appropriate environment could shape the effector function of MAIT cells.

Although human MAIT cells universally express RORγt and are abundant in both blood and tissues, only a small fraction – less than 5% in blood and 1% in the lung – produce IL-17^9,25^. The role of IL-23/IL-23R signalling, which drives Th17 differentiation and IL-17 secretion in conventional CD4^+^ T cells, has been investigated in MAIT cells but appears to be context dependent. In blood MAIT cells, single-cell studies show that IL-23 enhances cytotoxic function^27^, and dual stimulation through the TCR and cytokines is required to induce IL-17A production^25,26^. Cole *et al.* reported that IL-17 secretion by TCR+IL-12+IL-18 stimulation is independent of IL-23 signaling, whereas Wang *et al.* demonstrated that TCR+IL-23 stimulation can drive IL-17A producing MAIT cells to produce IFN-γ^25,26^. Our results not only confirm the findings from recent work, but also extend them by elucidating specific attributes of the IL-23 conditioning regimen that promote IL-17A production. We conclude that while IL-23R expression is necessary for IL-17A production, activation through this receptor must be complemented by TCR signaling. These requirements are best met in tissue environments, such as the lung, where both MR1-specific antigens and proinflammatory cytokines (IL-23 or IL-12+IL-18) are present.

Understanding how IL-23/IL-23R signaling shapes MAIT cell function and plasticity could have important implications for vaccine design. We postulate that adjuvants capable of enhancing IL-23 production or potentiating IL-23R signaling could result in augmented MAIT cell effector responses. For example, Wang *et al.* demonstrated that co-administration of IL-23 with 5-OP-RU improved control of pulmonary *Legionella* infection in mice and a 50-fold reduction in bacterial burden^29^. Moreover, Riffelmacher *et al.* showed that pulmonary immunization with an attenuated *Salmonella* strain expands two discrete MAIT cell lineages with distinct protective roles: an IL-17A-producing subset that protects against *Streptococcus pneumoniae,* and an IFN-γ-producing subset that protects against influenza^22^. Together with our findings, these studies suggest that targeted activation of IL-23R and the downstream STAT3 and type 17-associated genes can bias MAIT cells toward MAIT17 or mixed MAIT1/17 effector states, enhancing protective immunity against certain infections. Therefore, modulation of IL-23/IL-23R signaling is a potential therapeutic target to be considered when designing MAIT-targeted vaccines or immunotherapies.

## Method

### Study participants

#### Oregon Health & Science University, Oregon, USA

This study was conducted according to the Declaration of Helsinki. All samples and consent forms were collected, and all experiments were conducted according to protocols approved by the Institutional Review Board at Oregon Health & Science University (OHSU, IRB00000186). All ethical regulations relevant to human research participants were followed. Peripheral blood mononuclear cells (PBMCs) were obtained by apheresis from healthy adults. PBMCs were used for *in vitro* stimulation and to expand T cell clones as described below. Human serum was obtained from healthy adults.

#### Durban, South Africa

Bronchoalveolar lavage (BAL) fluid for isolating T cell clones and written informed consents from participants were collected according to a protocol approved by the University of KwaZulu Natal Human Biomedical Research Ethics Committee (UKZN BREC) and the Partners Institutional Review Board. Collection of excess fluid from adult patients undergoing clinically indicated diagnostic bronchoscopies was conducted at Inkosi Albert Luthuli Central Hospital. BAL-derived lung MAIT cell clones were isolated from participants, including patients with active tuberculosis and uninfected controls with suspected lung cancer as characterized in the previous study^9^.

### T cell clones

#### Isolation and characterization of T cell clones

For isolation of B1023-A5 and B1026-A8 clones, cells from BAL samples were stained with Propridium Iodide (Miltenyi Biotec), MR1/5-OP-RU tetramer (NIH tetramer core), and an antibody cocktail of surface stains to sort out non-CD3+ cells (PE/Cyanine7-conjuaged anti-CD14 (clone M5E2), anti-CD15 (clone HI98, anti-CD16 (clone B73.1), anti-CD19 (clone HIB19), anti-CD123 (clone 6H6), anti-CD235 (clone HIR2) antibodies from BioLegend). Live MR1/5-OP-RU tetramer positive cells were sorted and then cryopreserved on an Influx cytometer (BD Biosciences). After thawing, the cells were plated in a 96-well plate in a limited dilution assay with 1.5e5 cells irradiated PBMC (3000 cGray) and 3e4 cells irradiated LCL (6000 cGray) along with αCD3 (30 ng/mL), rhIL-2 (2 ng/mL), rhIL-12 (0.5 ng/mL; BioLegend), rhIL-7 (0.5 ng/mL; BioLegend), and rhIL-15 (0.5 ng/mL; BioLegend) in cytokine-supplemented RPMI 1640 medium (Gibco) containing 10% heat-inactivated human serum (R10HuS) in 96 well round bottom plates. On day 5, αCD3 was washed out by removal of half of the volume from the wells and replaced with additional cytokine-supplemented medium. Medium was changed for all wells every 2–3 days and replaced with cytokine-supplemented medium. T cell “buttons” were evaluated for growth on Day 20 and selected buttons stained with the MR1-5-OP-RU tetramer, PE/Cyanine7 anti-CD3 (clone SK7; BioLegend), FITC anti-CD4 (clone SK3; BioLegend), APC/Cyanine7 anti-CD8 (clone SK1; BioLegend), and BV605 anti-TCR Vα7.2 (clone IP26; BioLegend) antibodies. T cell clones that remained MR1-5-OP-RU tetramer positive were re-expanded in a 24 well non-ULA plate with irradiated 7.5e5 PBMC and 1.5e5 LCL as well as R10HuS supplemented with αCD3 and IL-2 but without additional rhIL-12, rhIL-7, or rhIL-15^31^. Briefly, human lung MAIT cell D1004-A3 clone and human lung non-MAIT clones (D0033-JE1) were previously isolated in the same manner as above, sorted for live MR1/5-OP-RU tetramer positive and live MR1/5-OP-RU tetramer negative CD8 positive cells, and characterized at the same time as other D1004-associated clones^9^.

#### Rapid expansion protocol

For expansion, T cell clones were co-cultured with a ratio of 5:1 for irradiated allogenic PBMCs and irradiated allogenic lymphoblastoid cell lines in R10HuS and αCD3 antibody (30 ng/mL)^38^. Recombinant interleukin-2 (2 ng/mL) was added the following day and every 2-3 days thereafter. T cell clones were washed on Day 5 to remove anti-CD3 antibody. Clones were frozen down at −80°C for 1-1.5e6 cells/mL after at least 11 days and used from frozen. New freezeback stocks were validated before use by flow cytometry.

### Reagents and Chemicals

5-(2-oxopropylideneamino)-6-d-ribitylaminouracil (5-OP-RU) was made from 5-A-RU*HCl that was prepared by the OHSU Medicinal Chemistry Core following a previously described procedure^39^. To prepare 5-OP-RU, equal volumes of circa 650 mM methylglyoxal (Sigma) and 32 mM 5-A-RU*HCl dissolved in water were combined immediately before addition to the assay to give a stock solution of 16 mM of 5-OP-RU^38^. Recombinant human IL-23 protein (R&D systems) was aliquoted and frozen at −20°C. IL-23 protein was diluted in R10HuS to appropriate conditions before use.

### Stimulation with IL-23 cytokine

#### IL-23 conditioning of PBMC

PBMC from a healthy donor were thawed at Day 0 and rested overnight in R10HuS. On Day 1 and 7, PBMC were stimulated with 5-OP-RU (1 µM). Starting at Day 1, PBMC were added with recombinant IL-2 (20 ng/mL) and with or without recombinant IL-23 (20 ng/mL) every other day until Day 11. Cells were harvested and analyzed for flow cytometry on Day 12.

#### IL-23 conditioning of MAIT cell clones

T cell clones were thawed and co-cultured with irradiated allogenic PBMCs and irradiated allogenic lymphoblastoid cell lines in R10HuS. On Day 0 and 6, T cell clones were stimulated with 5-OP-RU (1 µM). Starting at Day 0, T cell clones were added with recombinant IL-2 (2 ng/mL) and with or without recombinant IL-23 (20 ng/mL) every other day until Day 10. Cells were harvested and analyzed for flow cytometry assay or single-cell sequencing.

#### 1-day short-term IL-23 stimulation on MAIT cell clones for single-cell sequencing

5e5 T cell clones were plated and incubated with following conditions: no stimulation, 1-day stimulation with IL-23 (200 ng/mL), 3-hr PMA/Ionomycin stimulation. After stimulation, T cell clones were harvested and prepared for single-cell sequencing.

### Flow cytometry assays

The following reagents were obtained through NIH Tetramer Core Facility: MR1/5-OP-RU tetramer and MR1/6-FP tetramer conjugated to R-phycoerythrin (PE). Identification of MAIT cells with MR1 tetramers was described before^40^. To determine the total functional capacity of T cells, PBMC and T cell clones were incubated with PMA/Ionomycin (PMA, 20 ng/mL, Sigma; Ionomycin 1µM, Sigma) for 3 hours in the presence Brefeldin-A (5 µg/mL, BioLegend). For tetramer and intracellular cytokine staining, cells were then harvested and stained with MR1/5-OP-RU tetramer and MR1/6-FP tetramer at a 1:250 dilution in tetramer buffer (PBS containing 2% FBS) for 45 minutes at room temperature. After 45 minutes, cells were stained with Live/Dead Fixable Dead Cell Stain Ki (Thermo Fisher) and surface antibodies (APC anti-CD4 antibody, clone RPA-T4, BD Biosciences #555349; APC/Cyanine7 anti-CD8 antibody, clone SK1, BioLegend #344714; PE/Cyanine7 anti-TCR Vα7.2 antibody, clone 3C10, BioLegend #351711) for 30 minutes at 4°C. Cells were washed then fixed and permeabilized using Cytofix/Cytoperm (BD Biosciences) per manufacturer’s instructions. Cells were then stained FITC anti-IFN-γ antibody (clone B27, BioLegend #506504) and BV421 anti-IL-17A antibody (clone BL168, BioLegend #512322). All data was obtained with LSR II (BD) cytometer at the OHSU Flow Cytometry Shared Resource and analyzed with FlowJo software version 10 (TreeStar).

### RNA isolation, cDNA synthesis, and qPCR analysis

To measure *IL23R* expression, total RNA was isolated from human lung MAIT T cell clones (D1004-A3, B1023-A5, B1026-A8) using the RNeasy Plus Mini Kit (Qiagen). RNA was reverse transcribed into cDNA using the High-Capacity RNA-to-cDNA kit (Applied Biosystems) according to the manufacturer’s instructions. Quantitative RT-qPCR was performed on a Step One Plus Real-Time PCR System (Applied Biosystems) using TaqMan Universal PCR Master Mix (Life Technologies). Samples were run in triplicates and analyzed using Taqman probes (ThermoFisher Scientific) for *IL23R* (Hs00332759_m1) and for normalization *GAPDH* (Hs02758991_g1). Relative expression of human lung MAIT T cell clones were compared to human lung non-MAIT CD8 T cell clone (D0033-JE1).

### Single-cell sequencing of MAIT cell clones

After stimulation, T cell clones were incubated in Human TruStain FcX (BiolegendTM) for 10 minutes at 4°C and then stained with TotalSeq-C hashtag antibodies and CITE-seq antibodies (Biolegend^TM^) listed in Supplemental Table 3 for 30 minutes at 4°C. T cell clones were washed and counted to be loaded in 10X Genomics Chromium Next GEM Single Cell 5’ v2. Full protocol details of single-cell GEX and cell surface protein library preparation can be found from 10X genomics website (https://cdn.10xgenomics.com/image/upload/v1722286086/support-documents/CG000330_Chromium_Next_GEM_Single_Cell_5_v2_Cell_Surface_Protein_UserGuide_RevG.pdf). Finished libraries were then sent to Novogene Corporation Inc.TM in Sacramento California for NovaSeq 6000 sequencing.

### Single-cell RNA-seq pre-processing

Raw sequence reads were processed using 10X Genomics Cell Ranger software (version 6.1.1). The resulting sequence data were aligned to the GRCh38 human genome. Cell demultiplexing used a combination of algorithms, including GMM-demux, demuxEM and BFF, implemented using the cellhashR package^41–43^. Droplets identified as doublets (i.e. the collision of distinct sample barcodes) were removed from downstream analyses. We additionally performed doublet detection using DoubletFinder, and removed doublets from downstream analysis^44^. Next, droplets were filtered based on UMI count (allowing 0-20,000/cell), and unique features (allowing 200-5000/cell). Additionally, we computed a per-cell saturation statistic for both RNA and ADT data, defined as: 1 – (#UMIs / #Counts). This statistic provides a per-cell measurement of the completeness with which unique molecules are sampled per cell and has the benefit of being adaptable across diverse cell types. Data were filtered to require RNA saturation > 0.35. Analyses utilized the Seurat R package, version 4.2^45^. Using standardized methods implemented in the Seurat R package, counts and UMIs were normalized across cells, scaled per 10,000 bases, and converted to log scale using the ‘NormalizeData’ function. These values were then converted to z-scores using the ‘ScaleData’ command. Highly variable genes were selected using the ‘FindVariableGenes’ function with a dispersion cutoff of 0.5. Principal components were calculated for these selected genes and projected onto all other genes using the ‘RunPCA’ and ‘ProjectPCA’ commands. Clusters of similar cells were identified using the Louvain method for community detection, and UMAP projections were calculated. CITE-seq data were CLR-normalized by lane, meaning raw count data are subset per lane, CLR normalization performed (as implemented in the Seurat R package, using margin = 1), using all QC-passing cells/lane. Normalization per-lane was performed to reduce batch effects.

### Statistical Analysis

Statistical analysis was done in R studio for all differential gene expression of single-cell gene expression data. Wilcoxon test for statistical significance of gene expression between different groups. For all analysis we considered a 2-log threshold change of more than 0.5 and a Bonferroni-corrected p-value of < 0.05 to be significant. For histograms, data were graphed and analyzed with Prism 10 (GraphPad). Two-tailed paired Student’s t-test and one-way ANOVA Tukey’s multiple comparisons test was used for statistical analysis.

## Data availability statement

All R studio code used in the generation of this analysis is freely available and uploaded to github (https://github.com/kaindylan/IL23R-MR1T-Paper/tree/main). All raw data was deposited to dbgap (accessions number: phs003810.v1.p1). The authors declare that the data supporting the findings of this study are available within the paper and its supplementary information files. The source data underlying the graphs in the paper can be found in the Supplementary Data. The accession number for the atomic coordinates of CB964 A2 TCR-MR1-5-OP-RU, along with associated structure factors, have been deposited at the protein databank (www.rcsb.org) with accession code 9MS0. The raw data supporting the conclusions of this article will be made available by the authors, without undue reservation.

## Author contributions

The experiments presented were conceptualized by SK, DK, DAL, GS, and DML. SK, DK, GS, MC, EW, and SK performed experiments, as well as, analysis and interpretation of data. DAL and DML supervised the work. DAL, GS, MC, BB, GM, TR, and DML provided advice and technical expertise. All authors contributed to revising and reviewing the manuscript. All authors approved the final version of the manuscript.

## Acknowledgments

The research reported in this publication used computational infrastructure supported by the Office of Research Infrastructure Programs, Office of the Director, of the National Institutes of Health under Award Number S10OD034224. The content is solely the responsibility of the authors and does not necessarily represent the official views of the National Institutes of Health. We acknowledge the assistance of the Oregon Clinical & Translational Research Institute, which is supported by the National Center for Advancing Translational Sciences, National Institutes of Health, through Grant Award Number UL1TR002369, and contributions of the OHSU Medicinal Chemistry Core (RRID:SCR_019048). Analytical flow cytometry was performed in the OHSU Flow Cytometry Shared Resource (RRID:SCR_009974).

---

**Supplementary Figure 1.**
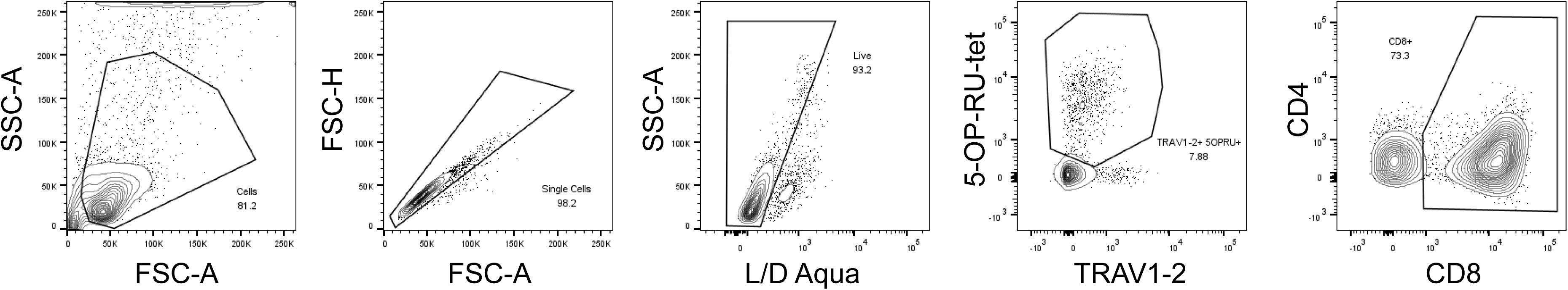
Gating strategy of MAIT cells from PBMCs. MAIT cells are defined by gating on cells, excluding doublets using forward scatter properties, selecting Live/Dead Aqua stain negative cells, followed by gating on TRAV1-2^+^, 5-OP-RU^+^, and CD8^+^ cells.

**Supplementary Figure 2.**
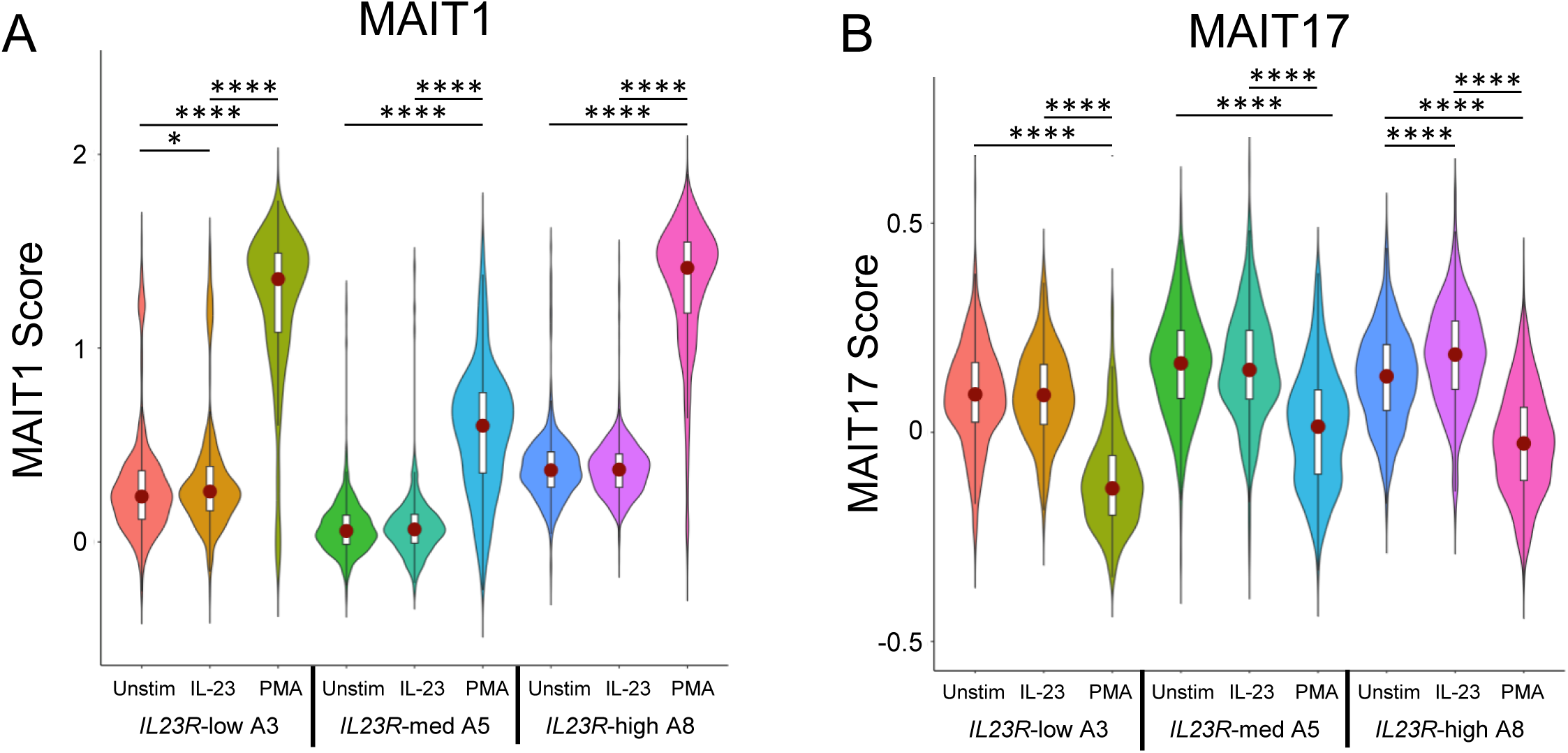
MAIT1 and MAIT17 transcriptional scores of MAIT cell clones. (A-B) MAIT1 (A) and MAIT17 (B) transcriptional scores of *IL23R*-low (A3), *IL23R*-med (A5), and *IL23R*-high (A8) for unstimulated, IL-23 stimulation, or PMA/Ionomycin stimulation. Kruskal-Wallis test. * = p ≤ 0.05, **** = p ≤ 0.0001.

**Supplementary Figure 3.**
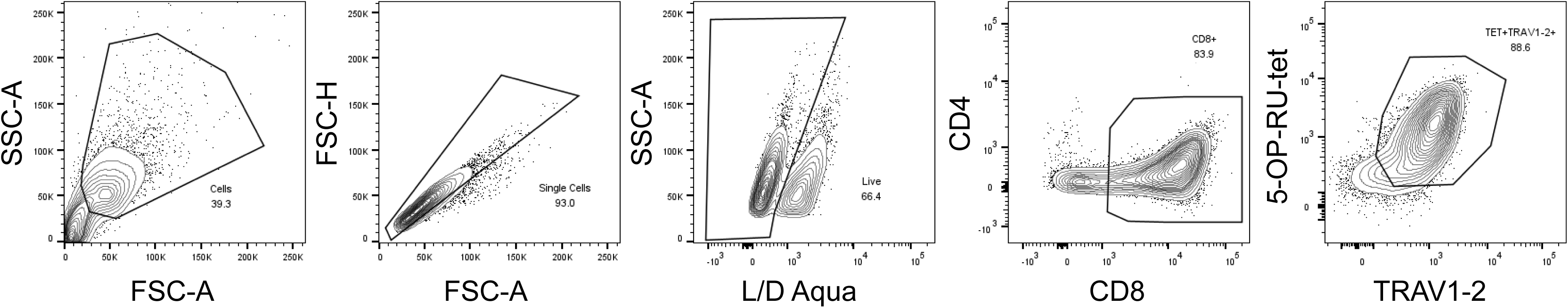
Gating strategy of MAIT cell clones. MAIT cells are defined by gating on cells, excluding doublets using forward scatter properties, selecting Live/Dead Aqua stain negative cells, followed by gating on CD8^+^, TRAV1-2^+^, and 5-OP-RU^+^ cells.

**Supplementary Figure 4.**
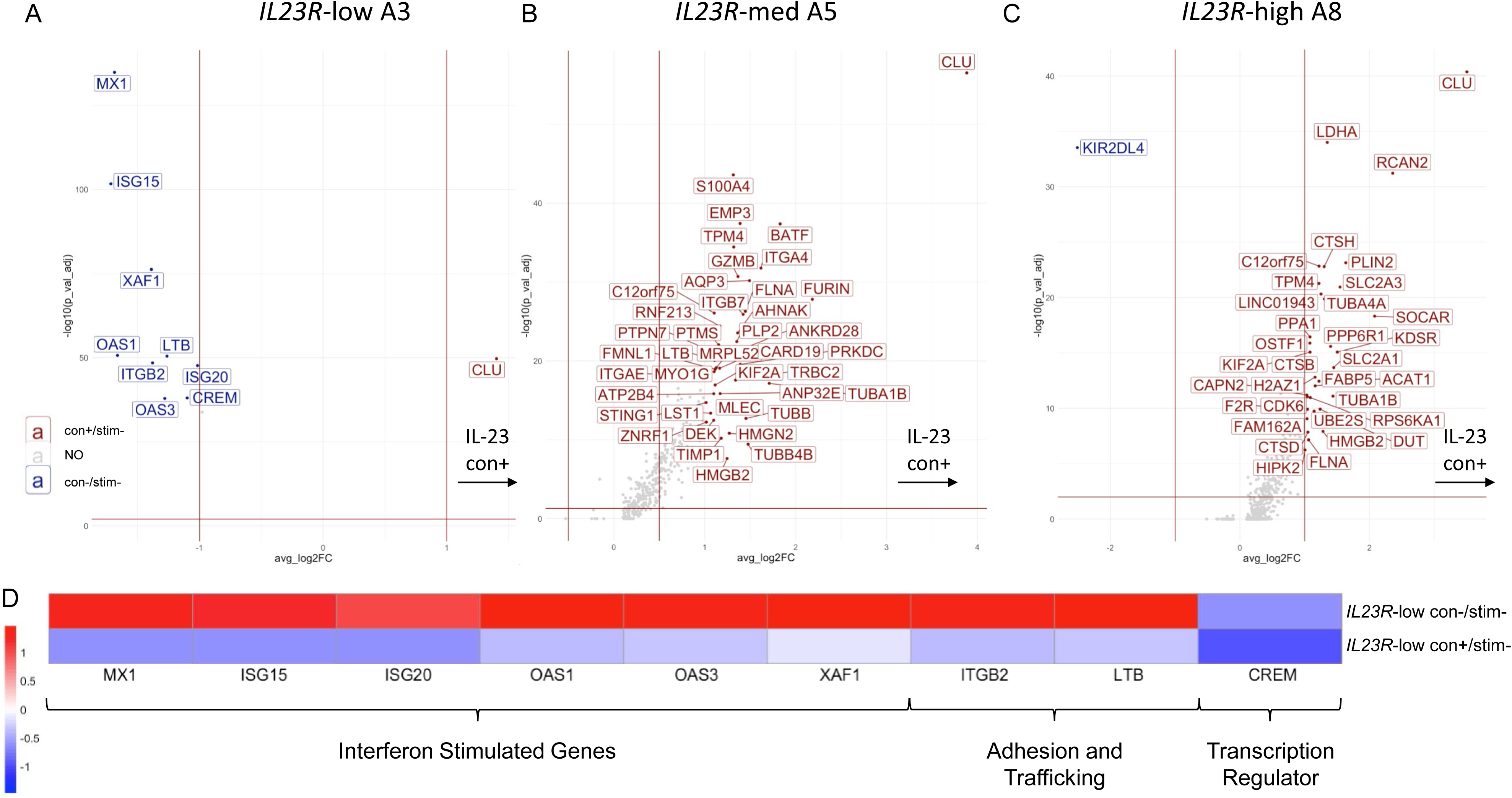
Differentially expressed genes with IL-23 conditioning. (A) Volcano plot of differentially expressed genes for D1004 A3 clone (low IL23R-expression). Red genes are significantly upregulated with IL23 conditioning and blue genes in unstimulated state. Genes expressed in at least 50% of cells, with at least 20% change between states, at least 1 log2 fold increased or decreased and p > 0.01 with Holm-Bonferroni correction. (B) Volcano plot of differentially expressed genes for B1023 A5 clone (medium IL23R-expression). Red genes are significantly upregulated with IL23 conditioning and blue genes in unstimulated state. Genes expressed in at least 50% of cells, with at least 20% change between states, at least 1 log2 fold increased or decreased and p > 0.01 with Holm-Bonferroni correction. (C) Volcano plot of differentially expressed genes for B1026 A8 clone (high IL23R-expression). Red genes are significantly upregulated with IL23 conditioning and blue genes in unstimulated state. Genes expressed in at least 50% of cells, with at least 20% change between states, at least 1 log2 fold increased or decreased and p > 0.01 with Holm-Bonferroni correction. (D) Heatmap of genes significantly downregulated in *IL23R*-low A3 with IL-23 conditioning. Genes are groups into shared categories and described.

**Supplementary Figure 5.**
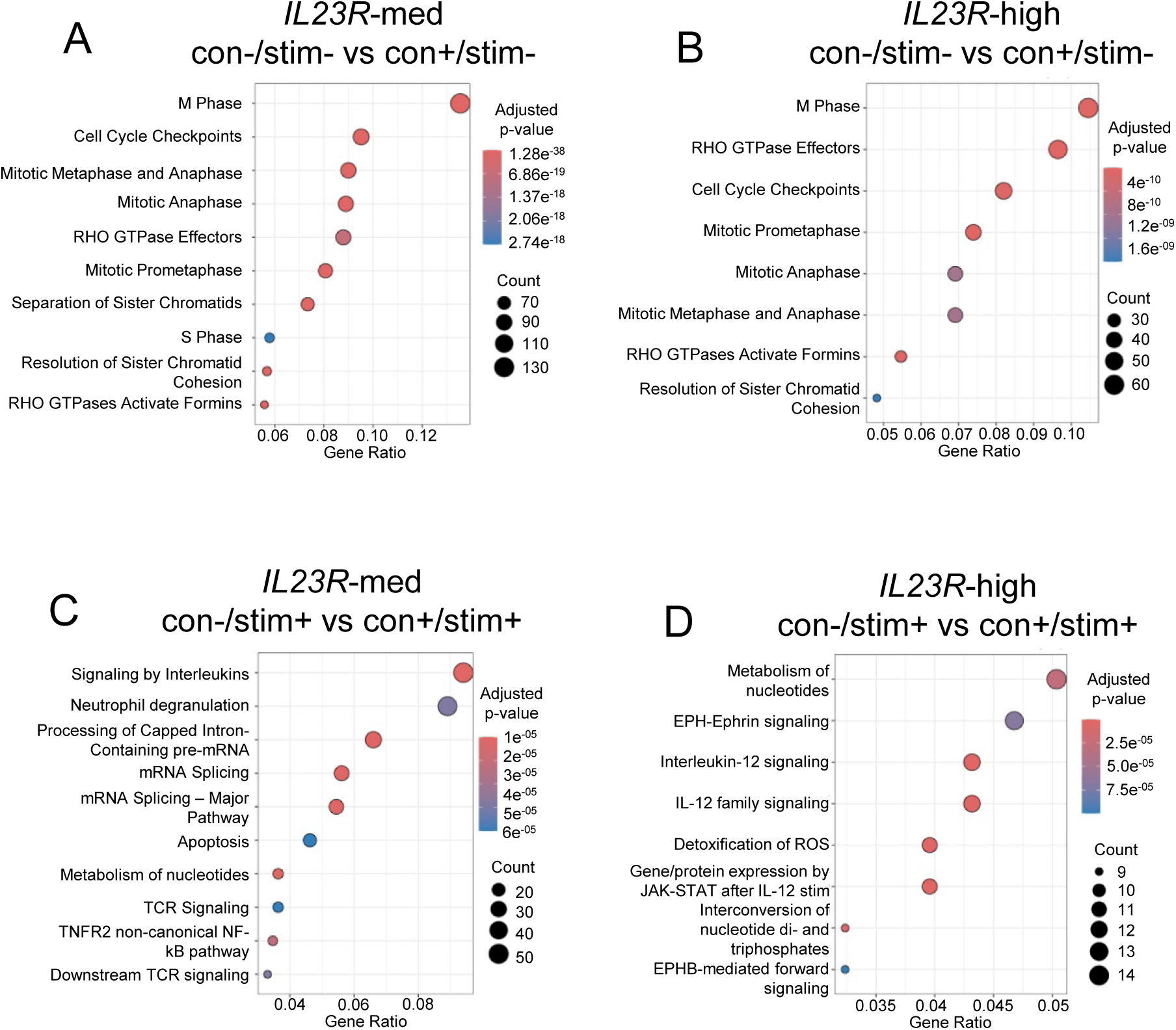
Pathway enrichment analysis of IL-23 conditioning and PMA/Ionomycin stimulation. (A-D) Pathway enrichment analysis for IL-23 conditioning versus no conditioning in *IL23R*-med (A5) (A) and IL23R-high (A8) (B), and for PMA/Ionomycin stimulation versus PMA/Ionomycin stimulation with IL-23 conditioning in *IL23R*-med (F) and IL23R-high (H) clones.

**Supplementary Figure 6.**
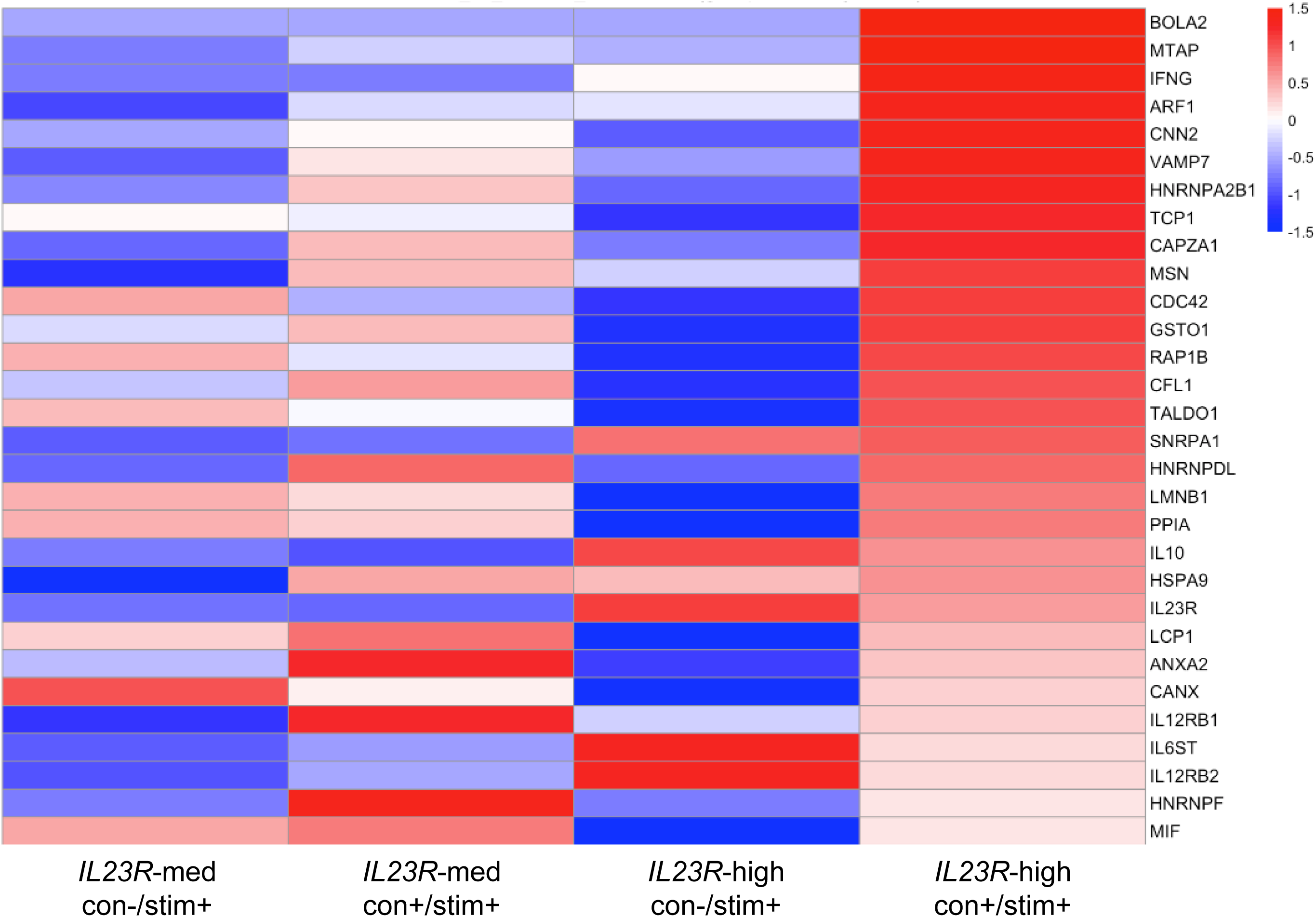
Heatmap of genes in IL-12 family signaling. Heatmap of genes upregulated in IL23R-high (A8) with IL-23 conditioning and PMA/Ionomycin stimulation as part of the IL-12 family signaling pathway.

**Supplemental Table 1. DE Results**

**Supplemental Table 2. Gene Comparison Matrix**

**Supplemental Table 3. Reagents used for 10X Genomics**® **Staining**

