## Supplemental Table 3 for "Antigenic stimulation in conjunction with cytokine is required for mediating IL-17A production in human MAIT cells"

Supplemental Table 3: Reagents used for 10X Genomics® Staining

| **Reagent** | **CAT** | **Company** |
| --- | --- | --- |
| TotalSeq Hashtag 1 | 394661 | Biolegend |
| TotalSeq Hashtag 2 | 394663 | Biolegend |
| TotalSeq Hashtag 3 | 394665 | Biolegend |
| TotalSeq Hashtag 4 | 394667 | Biolegend |
| TotalSeq Hashtag 5 | 394669 | Biolegend |
| TotalSeq CD4 | 300567 | Biolegend |
| TotalSeq CD8 | 344753 | Biolegend |
| TotalSeq CD26 | 302722 | Biolegend |
| TotalSeq CD161 | 339947 | Biolegend |
| TotalSeq CD197 (CCR7) | 353251 | Biolegend |
| TotalSeq CD279 (PD-1) | 329963 | Biolegend |
| TotalSeq CD45RA | 304163 | Biolegend |
| Human TruStain FcX | 422302 | Biolegend |
